# A Probabilistic Approach for Registration of Multi-Modal Spatial Transcriptomics Data

**DOI:** 10.1101/2021.10.05.463196

**Authors:** Yu Qiang, Shixu He, Renpeng Ding, Kailong Ma, Yong Hou, Yan Zhou, Karl Rohr

## Abstract

Observing the spatial characteristics of gene expression by image-based spatial transcriptomics technology allows studying gene activity across different cells and intracellular structures. We present a probabilistic approach for the registration and analysis of transcriptome images and immunostaining images. The method is based on particle filters and jointly exploits intensity information and image features. We applied our approach to synthetic data as well as real transcriptome images and immunostaining microscopy images of the mouse brain. It turns out that our approach accurately registers the multi-modal images and yields better results than a state-of-the-art method.

## 1. INTRODUCTION

Spatial transcriptomics (ST) technologies based on next generation sequencing (NGS) systematically generate spatial measurements of gene expression in an entire tissue sample, which bridge the gap between spatial information and the whole transcriptome [1]. Advanced spatial transcriptomics platforms, like Stereo-seq [2] or Seq-scope [3], achieve nanoscale resolution, enabling determining subcellular compartmentalization and visualization of RNA sequencing data.

However, for the current ST technologies with nanoscale resolution it is difficult to accurately assign spots of the transcriptome images (gene expression matrix images) to specific organelles or cells [1]. The information about gene expression can be exploited in different ways, for example, to characterize gene expression patterns or to classifiy cell types in the tissue [4–6]. However, lack of distinct cell boundaries in the transcriptome images presents a big challenge for automated analysis. Fast and accurate registration of transcriptome images and immunostaining images can facilitate the assignment of expressed genes to specific cells to enable studying sub-cellular gene expression patterns.

**Fig. 1.**
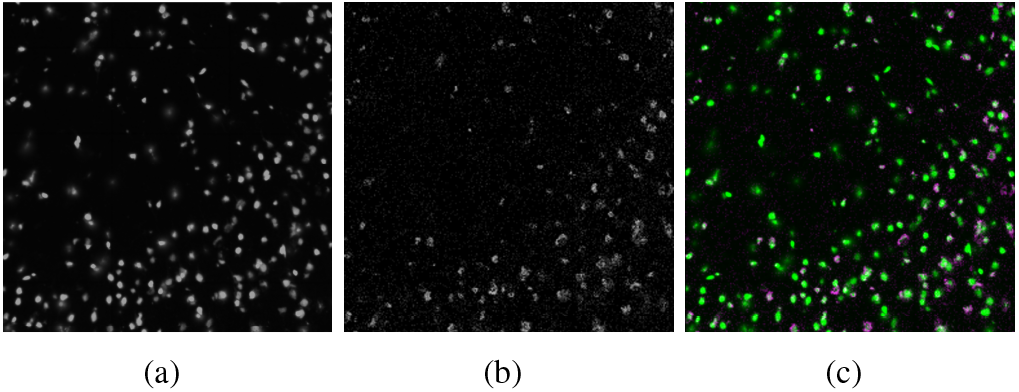
Registration result of our approach for an image section: (a) Immunostaining image, (b) transcriptome image (gene expression matrix image), and (c) overlay of registered images.

Although in situ sequencing (to generate transcriptome images) and immunostaining are carried out on the same tissue, there are multiple factors that can cause spatial shifting, for example, the sample preparation process and dispersion of RNAs after tissue permeabilization. Manual alignment of two images is time consuming, and only enables partial alignment in most cases [3, 6]. Therefore, methods are needed for efficient and accurate registration of large-scale transcriptome images and immunostaining images with tens of thousands of cells.

In previous work on automatic registration of spatial transcriptomics image data only few methods were introduced. [7] described a multiinformation-based method for registration of multiplexed in situ sequencing (ISS) datasets from the Human Cell Atlas project. Multiinformation is defined as KL divergence between a joint distribution and a product of marginal distributions, and used in conjunction with FRI (finite rate of innovation) sampling and swarm optimization. However, the method is computationally expensive since multiinformation is costly to compute.

In this work, we introduce a novel probabilistic approach for registration and analysis of transcriptome images and immunostaining images. The approach is based on a probabilistic Bayesian framework and uses particle filters to determine the transformation between the multi-modal images. Intensity information and image features are jointly taken into account.

We applied our approach to synthetic data as well as real transcriptome images and immunostaining microscopy images of the mouse brain. It turns out that our approach successfully registers the multi-modal images and yields better results than a state-of-the-art method.

## 2. METHODOLOGY

Our probabilistic registration approach consists of three main steps: (i) Generation of gene expression matrix image, (ii) spot detection and joint probabilistic registration, and (iii) Voronoi-based cell region determination.

### 2.1. Generation of expression matrix image

The gene expression matrix images (transcriptome images) are generated by inserting white dots on a black canvas (rectangular region for drawing dots). The x and y coordinates represent the horizontal and vertical positions of a pixel, respectively. In addition, the unique molecular identifier (UMI) count value of each position is used as the intensity value of the dot.

In order to emulate the real distance between dots, we adjust the dot size according to the relation between the dot size and the canvas image size. Let *d*_*spot*_ be the diameter of a circular dot, *w* the width of a canvas image (unit: inch), *h* the horizontal coordinate range of the gene expression matrix image, and *f*_*mag*_ the factor for dot magnification. The size of each dot is then calculated by:

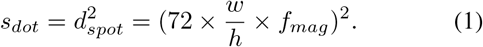

### 2.2. Probabilistic approach for registration

Our aim is to study the relationship between the spatial cell structure in the immunostaining image and the corresponding gene expression distribution in the gene expression matrix image (transcriptome image). The goal of registration is to assign *N*_*g*_ gene expression spots to *N*_*c*_ cells. Such assignment can be represented by a non-negative assignment matrix *ω* with elelments 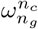 that denote the strength of gene expression for a spot *n*_*g*_ within a cell *n*_*c*_ (using a binary assignment 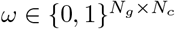). Some nodes may be assigned to no cells. In a Bayesian framework, we can formulate this task by estimating the posterior probability density *p*(*ω* | *I*^*i*^, *I*^*g*^), where *I*^*i*^ is the immunostaining image and *I*^*g*^ is the gene expression matrix image. We denote the positions of all detected spots in the immunostaining image by 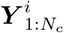 and the positions of all detected spots in gene expression image by 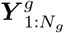 . To detect and localize multiple bright spots in an image, the spot-enhancing filter [8,9] is used. Since all cells are located within the same plane in our application, the transformation between the multi-modal images can be represented by a homography ***H***. Since 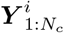 and 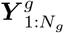 are conditionally independent of *I*^*i*^ and *I*^*g*^, by using Bayes’ theorem, we can write

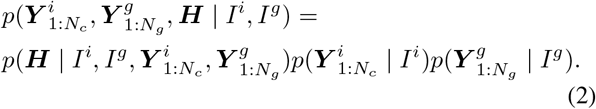

In our work, the transformation matrix is represented by

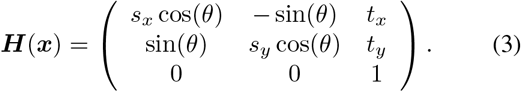

To determine ***H***, we need to compute the 5*D* parameter vector *x* = (*s*_*x*_, *s*_*y*_, *θ*, *t*_*x*_, *t*_*y*_) with rotation angle *θ*, scaling *s*_*x*_, *s*_*y*_, and translation *t*_*x*_, *t*_*y*_. As similarity metric between corresponding images we suggest using a combination of mutual information (MI) for the image intensities and the point set distance. The metric is maximized to align the multi-modal images.

Let MI (*I*^*i*^, ***H***(*x*)*I*^*g*^ be the mutual information of *I*^*i*^ and ***H***(*x*)*I*^*g*^, and D(***Y***^*i*^, ***H***(*x*)***Y***^*g*^) be the sum of distances to closest points (nearest neighbors) between points from ***Y***^*i*^ and ***H***(*x*)***Y***^*g*^. We search the optimal parameter vector *x*^∗^ by maximizing the following likelihood function

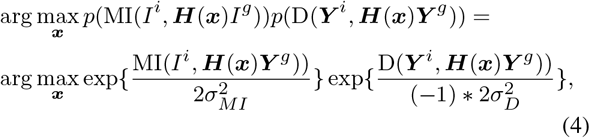

where *σ*_MI_ is the standard deviation of MI and *σ*_D_ is the standard deviation of the point set distance. Since MI(*r*) ≥ 0 and D(*r*) ≥ 0, the first term and the second term in Eq. (4) have opposite monotonicity.

We could naively search the whole parameter space to find the maximum likelihood to achieve the best alignment between the two images. However, this is time consuming. Here, we suggest using particle filters to efficiently determine the optimal *x*\* in Eq. (4), given a random initialization *x*.

Furthermore, we can estimate 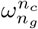 for each gene expression spot *n*_*g*_ at each cell *n*_*c*_ for a given observations 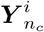 and 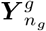. We use Voronoi tessellation [10] to identify cell regions (rough boundary of a cell) to assign gene expression spots to cells.

## 3. EXPERIMENTAL RESULTS

We compared our probabilistic registration approach with the state-of-the-art registration approach [7] using synthetic as well as real immunostaining images and gene expression matrix images. For synthetic data, we generated immunostaining images (500 × 500 pixels) using SimuCell [11] and randomly determined transformations as ground truth (GT) transformations. For the real data of the mouse brain (Stereo-seq [2], 964 × 964 pixels, pixel size 0.65 × 0.65*μ*m^2^), the GT transformation was obtained by manual alignment.

**Fig. 2.**
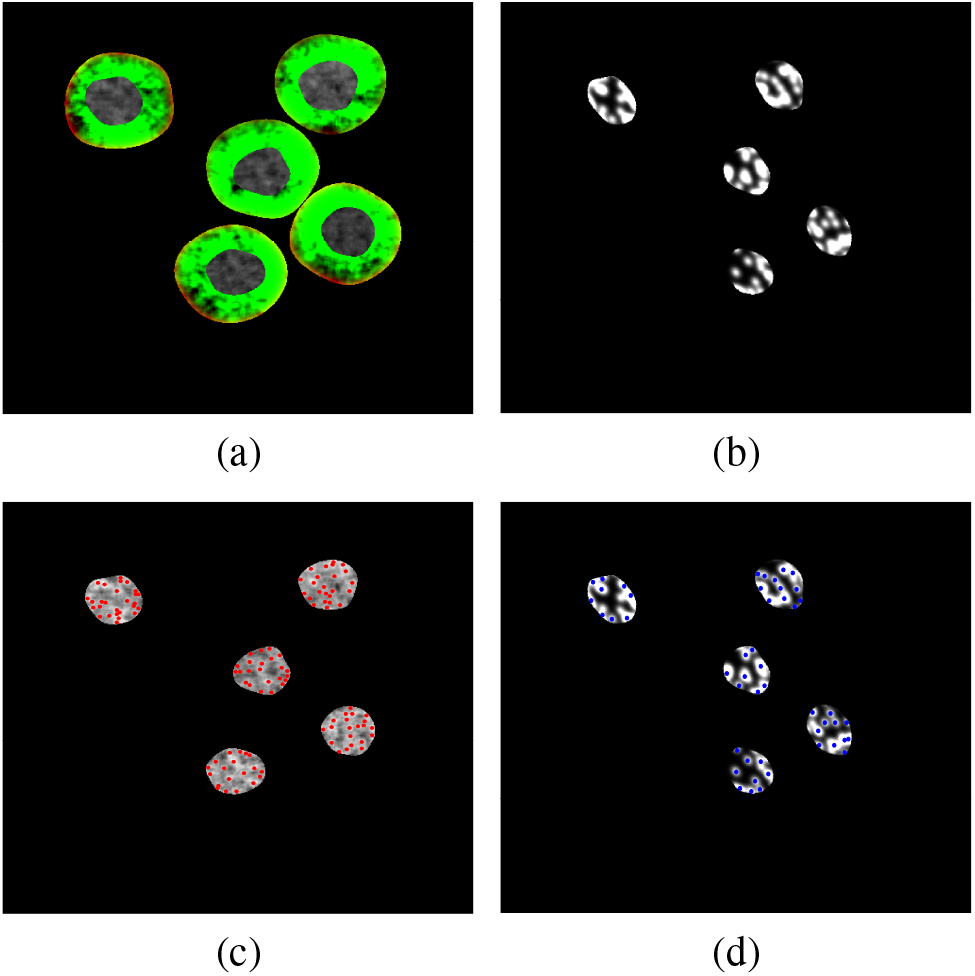
Results of our method for synthetic data: (a) Immunostaining image, (b) gene expression matrix image, (c) detected spots in immunostaining image (red), and (d) detected spots in gene expression matrix image (blue).

The target registration error (TRE) is used to quantify the registration result. The TRE is the average distance between the positions determined by the computed transformation parameters and the positions using the GT parameters. We employed six points to compute the TRE and determined the average position error in the *x* and *y* directions of the corresponding aligned points. Let ***H***(*x*\*) be the computed transformation matrix and *p*_*q*_, *q* ϵ {1, … , 6} be the manually selected points, then the TRE is computed by

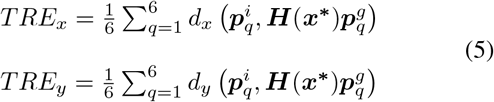

where *d*_*x*_, *d*_*y*_ denote the distance between two corresponding points in *x* and *y* direction. 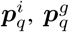 are two corresponding points in the immunostaining image and the gene expression matrix image.

For our probabilistic registration approach, we used 50 random samples and 10 time steps for the particle filter to determine the transformation matrix in Eq. (3).

**Fig. 3.**
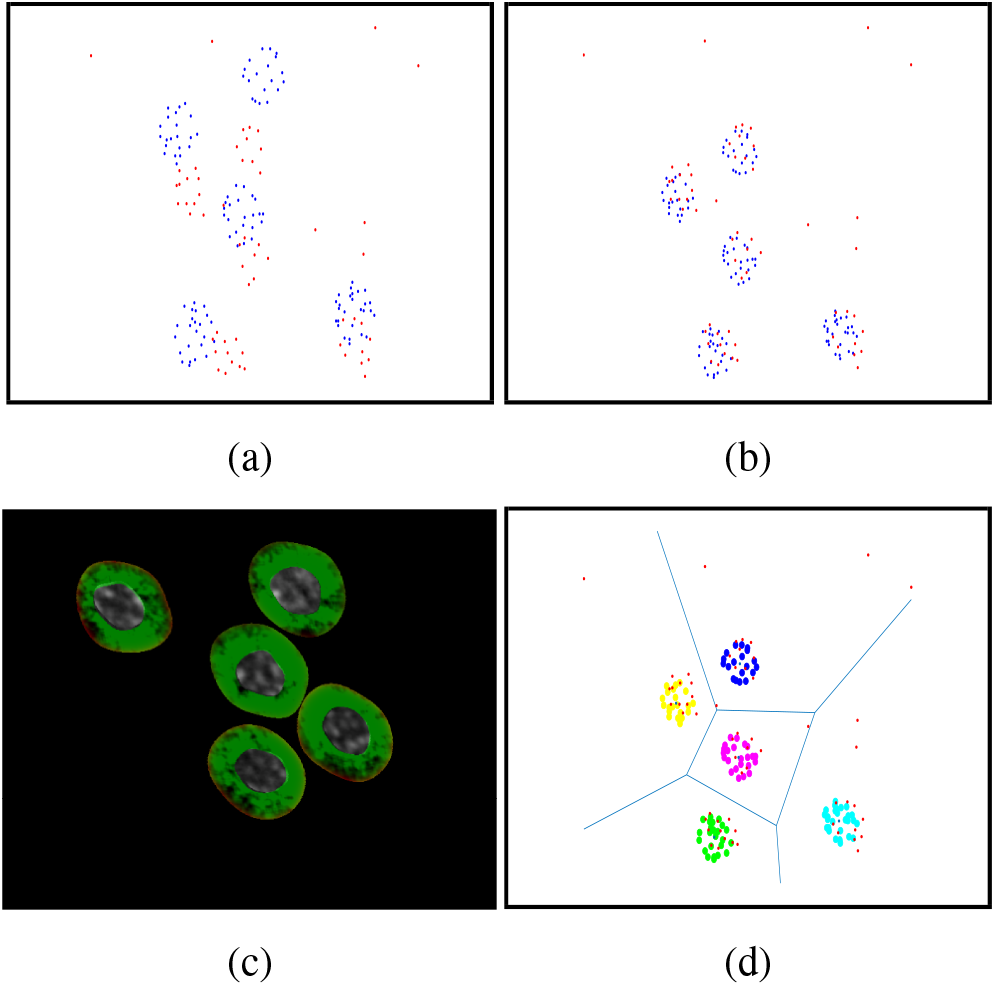
Results of our method for synthetic data with noise: (a) Overlay of detected spots in both images before registration (red: detected spots in immunostaining image, blue: detected spots in gene expression matrix image), (b) overlay of detected spots in both images after registration, (c) overlay of registered immunostaining image and gene expression matrix image, (d) computed cell regions by Voronoi tesselation (each cell is represented by a different color).

Table 1 and Table 2 provide the TRE of our registration approach and the method in [7] for four pairs of synthetic images and 4 pairs of real images. For all image pairs, the TRE of our approach is much lower than that of method [7]. Thus, our approach is much more accurate than the previous method for the considered challenging data. Further, the computational performance of our approach is about 3 times faster than the previous method [7].

**Table 1.** Target registration error for synthetic images.

| Image | #Cells | Method [7] |  | Ours |  |
| --- | --- | --- | --- | --- | --- |
| | | $TRE_x$ | $TRE_y$ | $TRE_x$ | $TRE_y$ |
| 1 | 5 | 8.32 | 15.65 | 2.27 | 9.78 |
| 2 | 20 | 15.14 | 17.37 | 2.88 | 13.37 |
| 3 | 80 | 6.84 | 20.74 | 3.29 | 15.38 |
| 4 | 320 | 12.89 | 23.84 | 5.17 | 17.16 |
| Average | 106 | 10.80 | 19.40 | 3.40 | 13.92 |

**Table 2.** Target registration error for real images.

| Images | #Cells | Method [7] |  | Ours |  |
| --- | --- | --- | --- | --- | --- |
| | | $TRE_x$ | $TRE_y$ | $TRE_x$ | $TRE_y$ |
| 1 | 256 | 22.68 | 23.17 | 12.21 | 16.56 |
| 2 | 153 | 18.14 | 15.37 | 8.44 | 15.42 |
| 3 | 175 | 22.84 | 27.74 | 10.29 | 13.38 |
| 4 | 208 | 23.89 | 22.84 | 10.17 | 14.16 |
| Average | 198 | 21.89 | 22.28 | 10.28 | 14.88 |

**Fig. 4.**
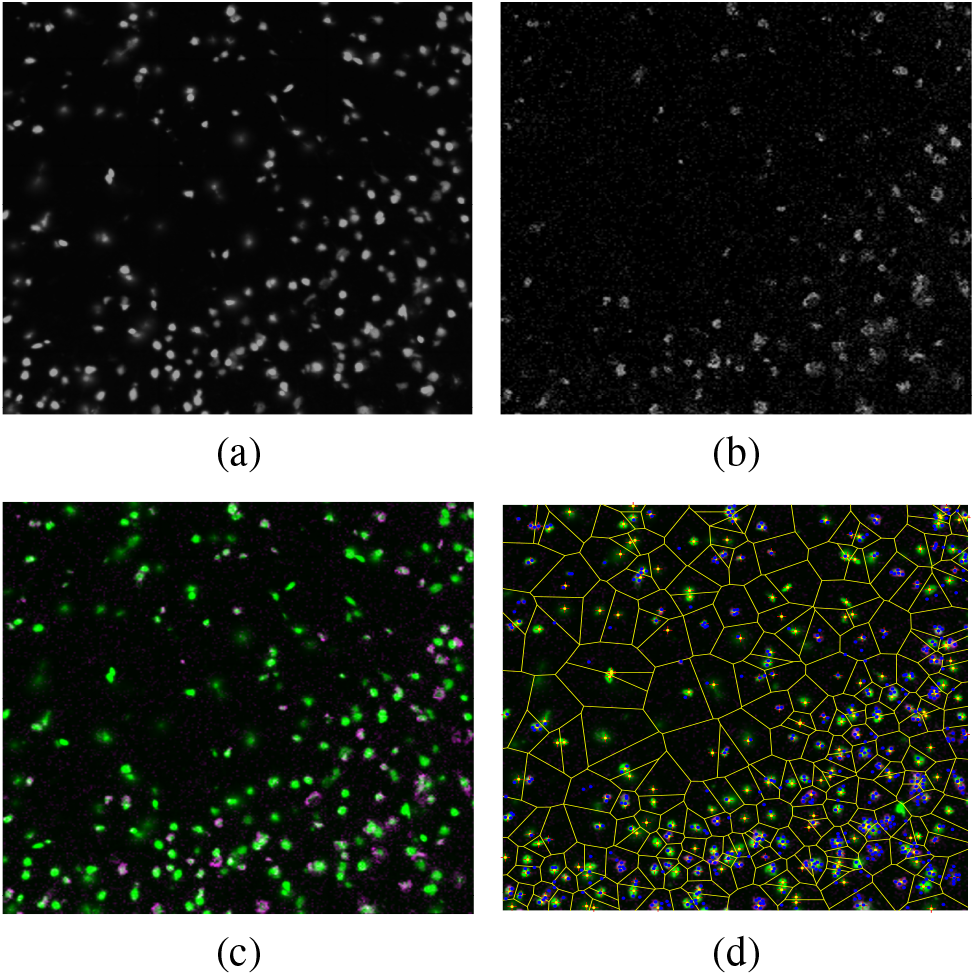
Results of our method for real data (image section): (a) Immunostaining image, (b) gene expression matrix image, (c) overlay of registered images, (d) computed cell regions (red crosses: detected center points of nuclei in immunostaining image using the spot-enhancing filter [8], blue points: detected spots in gene expression matrix image, yellow lines: computed cell regions).

## 4. CONCLUSIONS

We have presented a probabilistic approach for multi-modal registration of transcriptomics image data. Our approach determines the transformation between immunostaining images and gene expression matrix images. The method jointly exploits intensity information and image features. Our approach has been successfully applied to synthetic data and real spatial transcriptomics data of the mouse brain, and we found that our approach yields better results than a state-of-the-art method.

## 5. ACKNOWLEDGEMENTS

This work was supported by the BGI-Research and the Institute of Neuroscience (ION) of the Chinese Academy of Sciences. In addition, The authors gratefully acknowledge Prof Mu-ming Poo, Qing Xie, and Ao Chen for providing mouse brain spatial transcriptomics data and many helpful discussions during development.

## Compliance with Ethical Standards

This study was performed retrospectively using animal subject data from a previous study conducted under the approval of the Animal Care and Use Committee of Institute of Biomedicine and Health.

